# Metagenomes of Red Sea subpopulations challenge the use of morphology and marker genes to assess *Trichodesmium* diversity

**DOI:** 10.1101/2022.02.13.480231

**Authors:** Coco Koedooder, Etai Landou, Futing Zhang, Siyuan Wang, Subhajit Basu, Ilana Berman-Frank, Yeala Shaked, Maxim Rubin-Blum

## Abstract

The bloom forming *Trichodesmium* are filamentous cyanobacteria of key interest due to their ability to fix carbon and nitrogen within an oligotrophic marine environment. *Trichodesmium* blooms typically comprise a complex assemblage of subpopulations and colony-morphologies that are predicted to exhibit distinct ecological lifestyles. Here, we assessed the poorly studied diversity of *Trichodesmium* in the Red Sea, based on metagenome-assembled genomes (MAGs) and *hetR* gene-based phylotyping.

We assembled four non-redundant MAGs from morphologically distinct *Trichodesmium* colonies (tufts, dense and thin puffs). *T. thiebautii* (puffs) and *T. erythraeum* (tufts) were the dominant species within these morphotypes. While subspecies diversity is present for both *T. thiebautii* and *T. erythraeum*, a single *T. thiebautii* genotype comprised both thin and dense puff morphotypes, and we therefore hypothesize that the phenotypic variation between these morphologies is likely attributed to gene regulation. Additionally, we found the rare non-diazotrophic clade IV and V genotypes, related to *T. nobis* and *T. miru* respectively, that likely occurred as single filaments. *HetR* gene phylogeny indicates that the genotype in clade IV could represent the species *T. contortum*.

We further show that *hetR* phylotyping can overestimate the taxonomic diversity of *Trichodesmium*, as two copies of the *hetR* gene were present within *T. thiebautii* genomes, one of which misidentified this lineage as *T. aureum*. Taken together, our results highlight the importance of re-assessing *Trichodesmium* taxonomy while showing the ability of genomics to capture the complex diversity and distribution of *Trichodesmium* populations.

## Introduction

*Trichodesmium* is a genus of filamentous cyanobacteria known for its ability to form large visible surface blooms in tropical and subtropical regions of the ocean. Having first been described in 1770 by James Cook^1^, *Trichodesmium* have been extensively studied due to their ability to form blooms of large biomass supporting marine food webs through their nitrogen and carbon fixing capabilities^2^ and subsequent relevance to biogeochemical cycles within oligotrophic marine environments^3–5^.

*Trichodesmium* blooms are dynamic over space and time, typically consisting of a complex assemblage of several different subpopulations and colony morphologies that are predicted to exhibit unique ecological lifestyles^6–11^. For example, the comparison of two distinct puff-shaped morphotypes termed ‘dense’ and ‘thin’ colonies, isolated from the Red Sea displayed a remarkable heterogeneity in their preference to capture and center dust^7^. While colony morphotypes cannot always be linked to different genotypes or species^8,12^ genomic information regarding *Trichodesmium* colonies in this region is lacking in comparison to studies conducted in the Atlantic and Pacific Oceans.

The diversity of *Trichodesmium* bloom-forming populations was first assessed via colony morphology^13^. With the advent of molecular techniques, the use of single-marker genes represented a more accurate and consistent phylogenetic technique^14^. Marker genes that have been used to assess *Trichodesmium* diversity include the 16S rRNA, *hetR*, a regulatory gene for the development of heterocysts, and *nifH*, which encodes the nitrogenase iron protein. The *hetR* gene is considered a good marker to assess *Trichodesmium* diversity as it is more variable (10%) in comparison to genetic markers such as 16S ribosomal RNA (2-3%) or *nifH* (2%)^15,16^.

Phylogenetic analysis using *hetR* as a marker gene separates *Trichodesmium* into four distinct clades: clade I (*T. thiebautii*, *T. hildebrandtii*, *T. tenue*, *T. pelagicum*), clade II (*T. aureum*), clade III (*T. erythraeum*, *T. havanum*), and clade IV (*T. contortum*, *T. tenue*)^15,17^. Recently, a fifth clade (clade V) consisting of the non-diazotrophic *T. miru* was proposed by Delmont (2021). Currently, genomes of clades I, III, (and V) are available, including that of the culturable clade III lineage *Trichodesmium erythraeum* IMS101^18^.

To better understand the taxonomic diversity of *Trichodesmium* in the Red Sea, we isolated colonies from the Gulf of Aqaba and compared the resulting metagenome-assembled genomes (MAGs) to those of other *Trichodesmium* populations from the Indian, Pacific and Atlantic Oceans. We confirmed the presence of these genomes in the Red Sea based on amplicon sequencing of the *hetR* gene, and, in light of our findings, evaluated the ability of this marker gene to capture *Trichodesmium* diversity.

## Material and Methods

### Trichodesmium sampling and extraction

To assess the *Trichodesmium* population of the Red Sea, *Trichodesmium* colonies were handpicked and separated into three distinct morphotypes (~100-200 colonies each) throughout the winter bloom (November 2020) using a 100 μm phytoplankton net at 20 m depth in the Gulf of Aqaba (Eilat, Israel) (29.56°N, 34.95°E) (Figure 1). Colonies were washed 3 times in 0.2 μm filtered seawater before being filtered on a 0.2 μm PCC filter using a vacuum pump. Filters were frozen in liquid N2 and kept at −80°C until analysis.

**Figure 1.**
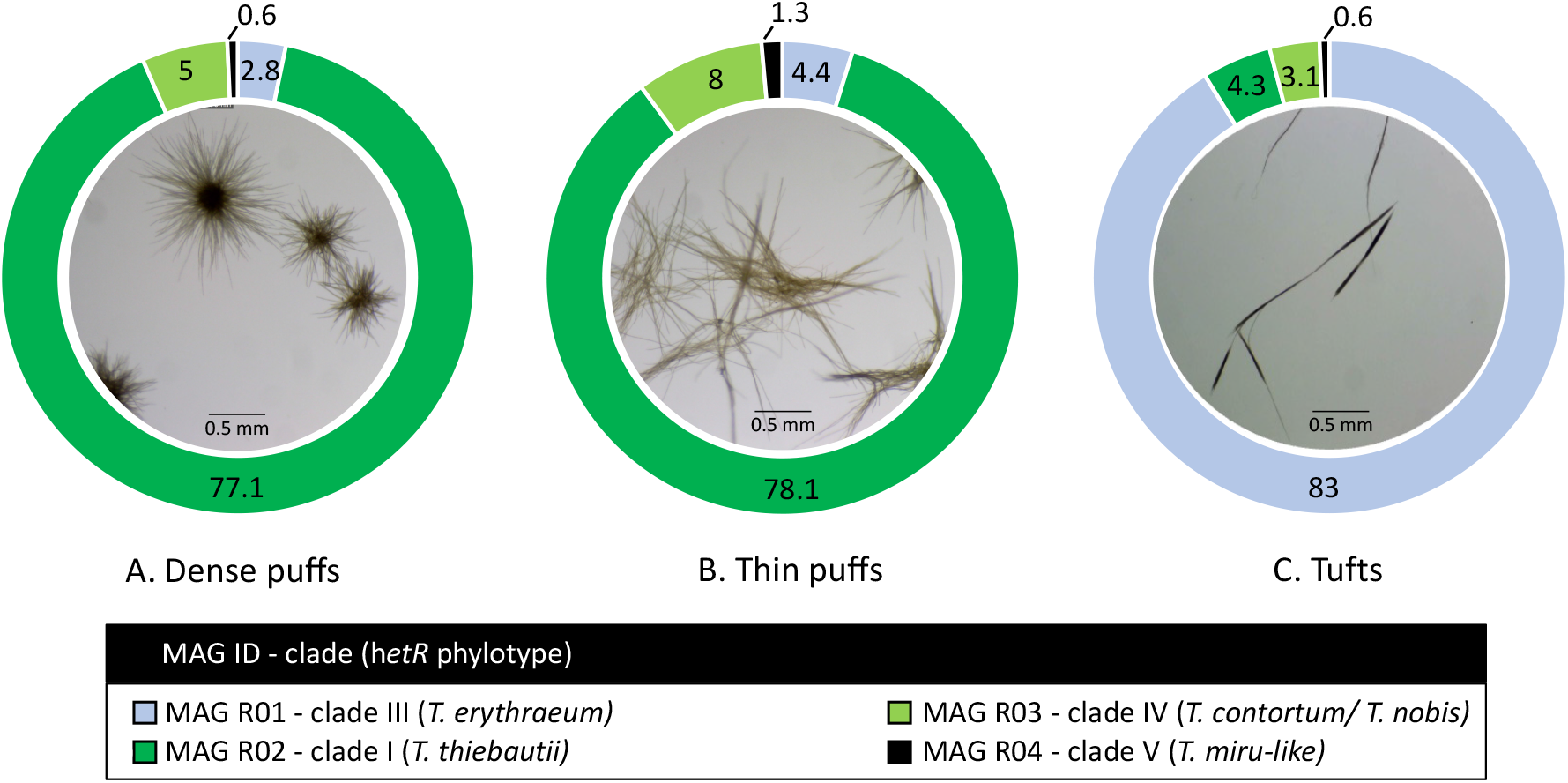
*Trichodesmium* colony morphotypes and their relative read abundance (%) in the Red Sea. Both **(a)** dense and **(b)** thin puff-shaped morphotypes, primarily consisted of *T. thiebautii* genotype (dark green; MAG R02), while tuft-shaped morphotypes **(c)** were dominated by *T. erythraeum* (light blue; MAG R01). All morphotypes contained low read abundance of non-diazotrophic MAG R03 (light green; MAG R03) and MAG R04 (black; MAG R04) genotypes, clustering within clade IV and V respectively.

DNA was extracted from three *Trichodesmium* samples using the DNAeasy Plant Mini Kit (Qiagen) following the manufacturer’s instructions. Metagenomic libraries were prepared at HyLabs (Rehovot, Israel) and sequenced with 30-60 million 2 x 150 bp reads on Illumina HiSeq at GENEWIZ (Leipzig, Germany). The metagenomic samples can be found under the accession numbers SRR17940154-6.

### Metagenomic analysis

*Trichodesmium* metagenome-assembled genomes (MAGs) were binned from assemblies based on the raw reads from this study (PRJNA804487) and those from the previously published metagenomes of *Trichodesmium* from the Pacific (PRJNA435427; PRJNA358796) and Atlantic (PRJNA330990) Oceans^8,19,20^. Metagenomes were assembled, annotated, and binned using the ATLAS (v2) pipeline^21^ (Supplementary R Markdown File). Briefly, raw sequences underwent quality control through the BBTools suite^22,23^ and were assembled using metaSPAdes^24^ (k-mer lengths: 21, 33, 55, 99, 121 bp). MAGs were binned from each sample using MetaBAT 2^25^, MaxBin 2.0^26^ and VAMB^27^. The completeness and redundancy of each bin were assessed using CheckM^28^. A non-redundant set of bins was produced using DAS Tool^29^ and dRep^30^ based on an average nucleotide identity (ANI) cutoff of 97.5%. MAGs were taxonomically characterized using the genome taxonomy database tool kit GTDB-tk^31^. Genes were predicted using Prodigal^32^. We refer to the Red-Sea MAGs from this study as MAG R and MAGs based on other studies as MAG T. The genome of the cultured strain *Trichodesmium erythraeum* IMS101 and five TARA Oceans MAGs, including two novel non-diazotrophic genomes^6^, were incorporated in the follow-up analysis. The ANI values for each MAG were analyzed using ANIb module in pyANI^33^.

### Amplicon sequencing of the hetR gene

*Trichodesmium* samples (tuft and puff morphotypes) for amplicon sequencing of the *hetR* were collected in the Gulf of Aqaba (same location as above) during several blooms (2013-2019). DNA was extracted using the phenol-chloroform method^34^. Partial *hetR* gene sequences (~355 bp) were amplified using the forward primer (hetrR_50F), 5’-ATTGAACCYAAACGGGTT-3’ and reverse primer (hetR_381R), 5’ CGCTTAATATGTYCTGYCAAAGCTT-3’, which were deduced from conserved regions of a *Trichodesmium hetR* nucleotide sequence alignment. 2 x 250 bp reads were sequenced on Illumina MiSeq, following library preparation at HyLabs (Rehovot, Israel). The *hetR* amplicion samples can be found under the accession numbers SAMN25885796-803. Sequences were merged, denoised and called into amplicon sequence variants (ASVs) using DADA2 in Qiime2^35,36^. The ASVs were clustered into five OTUs at 98% similarity for further downstream analyses using the VSEARCH consensus taxonomy classifier ^37^.

### Trichodesmium phylogeny

We constructed multi-locus and single marker gene (*hetR*) phylogenies of *Trichodesmium*. We used the following genomes: the four *Trichodesmium* MAGs from this study, five *Trichodesmium* MAGs from the TARA Oceans dataset^6^ and that of *Trichodesmium erythraeum* IMS101 PRJNA318^18^. The phylogenomic tree was constructed from a concatenated gene-alignment of a 251 single-copy gene-set hidden Markov Models (HMMs) for Cyanobacteria using GToTree (v.1.16.12; default settings)^38^. The aligned protein sequences were refined using Gblocks (0.91b; default settings) to eliminate poorly aligned positions and divergent regions^39^. A tree was subsequently constructed from the cleaned alignment using IQtree2 with ModelFinder to estimate the best-fit model (Q.plant+F+I+G4)^40^. Shimodaira–Hasegawa approximate likelihood-ratio test (SH-aLRT) and ultrafast bootstrap approximation (UFBoot) branch support values were estimated from 1000 bootstraps. The tree was visualized using FigTree (v1.4.4) and rooted with *Okeania hirsuta* (GCA_003838225)^41^ as an outgroup (Supplementary Table 4).

The *hetR* marker gene was identified in *Trichodesmium* MAGs using the Rapid Annotation using Subsystem Technology (RAST)^42^. The *hetR* sequence and the neighboring genes cluster were identified by searching for the HetR amino acid sequence (Tery_1921; Q93CE9) using BLAST on the SEED server^43^. The marker gene sequences from each *Trichodesmium* MAG were aligned with previously published ones^16,17,44^ and those from our Red Sea *hetR* amplicon sequencing using the Multiple Alignment Fast Fourier Transformation (MAFFT; L-INS-i) software^45^. All the *hetR* sequences used in this study can be found in Supplementary Table 5. This alignment was further cleaned using Gblocks (0.91b; default settings; Castresana, 2000). A phylogenetic tree rooted at the sequence of *Okeania hirsuta*, was constructed using IQtree2 for *hetR* (best-fit model TPM3+F+I) and visualized using FigTree (Figure 3).

**Figure 2.**
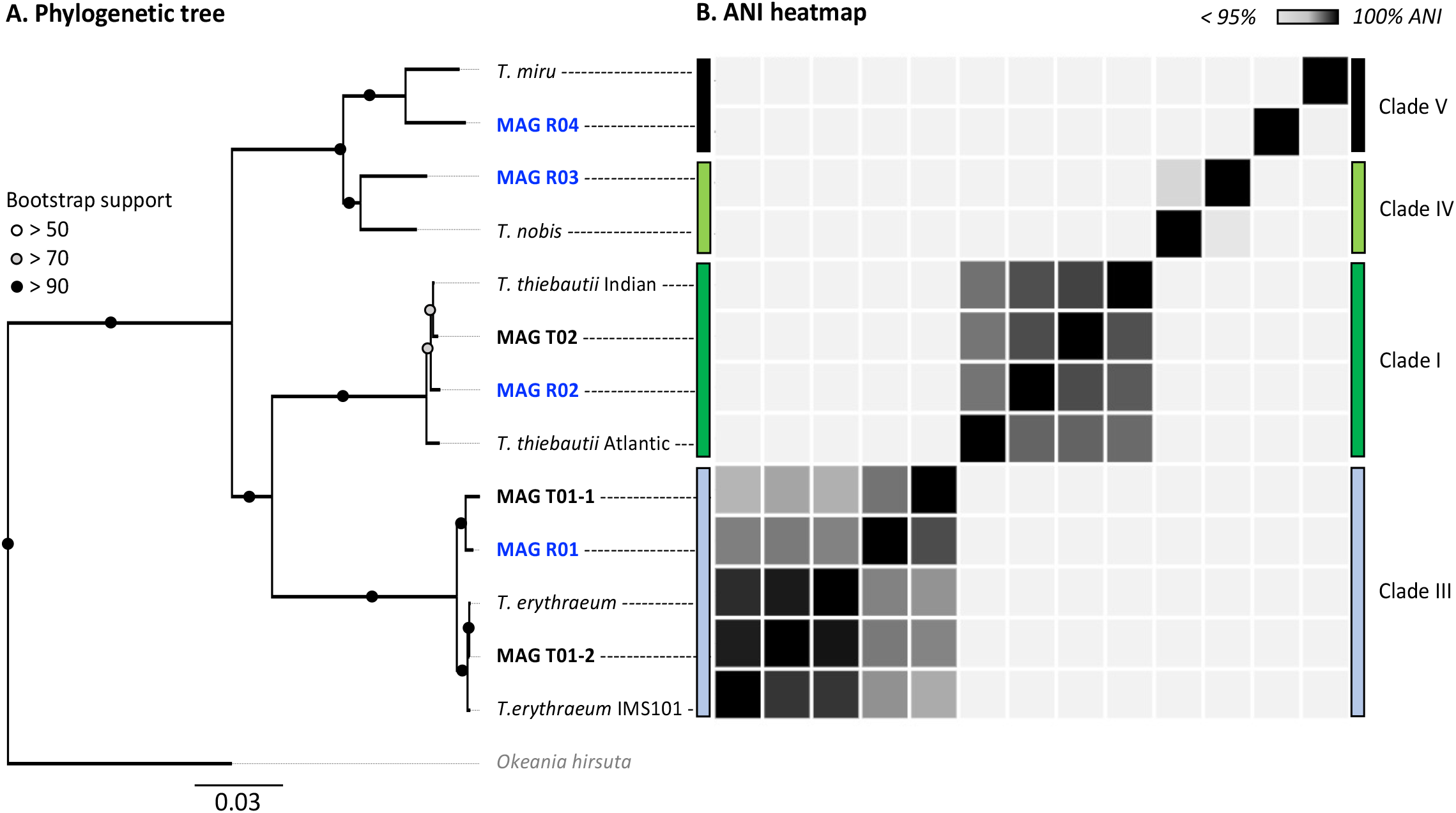
Multi-locus (251 HMMs) phylogenomic tree (a) and average nucleotide identity (ANI) heatmap (b) of *Trichodesmium* MAGs. All MAGs assembled in this study are marked in bold. Red Sea MAGs (our samples) are marked in blue. The phylogeny includes five MAGs from the TARA Oceans dataset and the laboratory culture *T. erythraeum* IMS101. The tree was rooted at *Okeania hirsuta* (grey). ANI values can be found in Supplementary Table 2.

**Figure 3.**
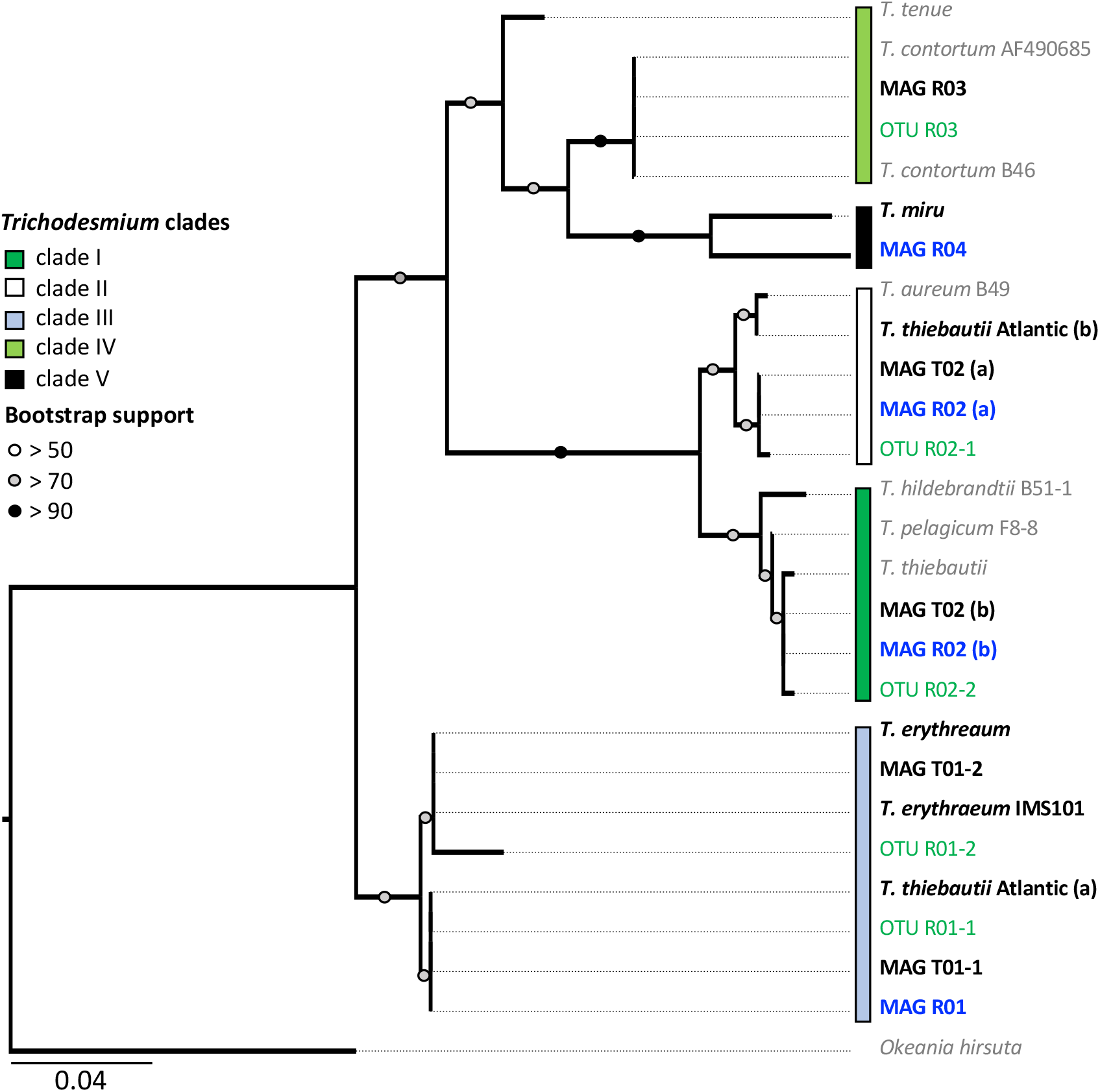
*Trichodesmium hetR* phylogeny. The tree is based on the alignment of 312 nucleotide sequences. Bold text indicates the *hetR* sequences from MAGs (Red Sea sequences are marked in blue). Amplicon *hetR* sequences are shown in green (Supplementary Table 4). The five proposed *Trichodemisum* clades are shown with colored boxes. Note that clades IV and V are polyphyletic and are therefore not completely resolved. MAGs R02 and T03 contain two *hetR* sequences, clustering within clades I and II, respectively. The *hetR* sequences from *T. erythraeum* genomes clustered into two groups within clade III. The percentage identity matrix between the *hetR* gene sequences can be found in Supplementary Table 3.

## Results and Discussion

Four non-redundant *Trichodesmium* MAGs were assembled from the Red Sea (Figure 1; Supplementary Table 1). The MAGs were of high quality, being 90% complete and with less than 5% redundancy, apart from MAG R04 which was 86.29% complete. To provide a wider framework of *Trichodesmium* diversity, we assembled three non-redundant *Trichodesmium* MAGs from the Pacific and the Atlantic Ocean, based on previously published raw data^8,20,46^. These included two high-quality MAGs T01-2 and T02, as well as the lower quality MAG T01-1 (Supplementary Table 2).

Our results suggest that *T. erythraeum* and *T. thiebautii* are the dominant *Trichodesmium* lineages in the Red Sea, where MAG R01 clustered with *T. erythraeum* (97.3% ANI), and MAG R02 with *T. thiebautii* (97.7-98.3% ANI) (Figure 2; Supplementary Table 3). The dominance of these lineages in the Red Sea could be confirmed from *hetR* gene phylotyping of *Trichodesmium* populations across several seasons (Supplementary Figure 1). Tuft colonies were primarily composed of *T. erythraeum* (~83% read abundance) (Figure 1c)^16^.

Both thin and dense puff-shaped colonies were dominated by *T. thiebautii* (~78% read abundance) (Table 1; Figure 1a, b). This finding is similar to another genomic study where radial (dense-like) and non-radial puff morphologies were linked to the same genotype of *T. thiebautii*^8^. Puff-shaped *Trichodesmium* colonies have the unique capability to interact, capture and concentrate dust at the colony’s core^47^ which can allow colonies to obtain limiting nutrients from the marine environment, such as iron or phosphorous^48,49^. In the field, thin puffs isolated from the Red Sea were shown to be more interactive in comparison to dense puffs in the capturing and concentrating of dust at the colony’s core, whilst exhibiting pronounced gliding motility^7^. Our findings highlight that these morphological variations do not appear to be explained genomically and that other yet uncharacterized factors are at play. We hypothesize that the observed variations in puff morphologies concerning the centering of dust^7,50^ are due to differences in gene regulation in response to yet unknown environmental conditions which may include nutrient limitation or grazing. In a laboratory setting, single filaments of *T. erythraeum* IMS101 clustered into the puff and tuft-shaped colonies when subjected to iron or phosphorous limitation^51^. Nonetheless, *Trichodesmium* displays functional variability and heterogeneity at a single-colony level that remains largely unexplained ^9,10^ and subsequently the ability to predict whether a colony will center dust still requires ongoing exploration.

Our data revealed subspecies diversity for both *T. thiebautii* and *T. erythraeum* (Figure 2). *T. thiebautii* MAGs appeared to cluster according to broad geographical patterns as MAG R02 was more similar to that of *T. thiebautii* populations isolated from the Indian Ocean (98.3% ANI), than to the Atlantic *T. thiebautii* MAG (97.7% ANI). This concurs with the previous findings^6^. We also found genomic diversity within *T. erythraeum* as revealed from two distinct *T. erythraeum* MAGs from the Pacific Ocean (96.6% ANI). MAG T01-1 was highly similar to that of *T. erythraeum* isolated from the Red Sea (97.6% ANI; MAG R01), whereas MAG T01-2 was closely related to *T. erythraeum* IMS101 (99.4% ANI), which was isolated from the Atlantic Ocean^52^. The *hetR* gene phylotyping of *Trichodesmium* populations in the Red Sea indicates the occurrence of an additional *T. erythraeum* subspecies similar to that of MAG T01-2 (Figure 3). We were, however, unable to assemble its genome or link it to a specific morphotype. It is still unclear if these two *T. erythraeum* subspecies within the Red Sea and the Pacific Ocean occupy distinct ecological niches.

We detected two potentially non-diazotrophic species within the Red Sea with minor read abundance in all three samples (Supplementary Table 1). Both genotypes could not be linked to a specific morphotype, and it is plausible that they occurred as single filaments within our samples. The low coverage reflected a previous estimate of the non-diazotrophic *Trichodesmium* of the Red Sea^6^. Whereas MAG R03 clustered within clade IV, which includes the non-diazotrophic *T. nobis* (95.6% ANI), and MAG R04 clustered within clade V containing the non-diazotrophic *T. miru* (94.6% ANI), both MAGs are on the threshold of being a distinct species within clade IV and V, respectively. We, therefore, turned to the single-marker gene *hetR* to further differentiate these MAGs, as *hetR* phylogeny captures a larger diversity of *Trichodesmium* than is possible through the limited number of *Trichodesmium* genomes currently available. The *hetR* sequences, but not the genomes, are available for *T. pelagicum* (AF490696.1), *T. hildebrandtii* (AF490679.1) in clade I, *T. aureum* (AF490680.1) in clade II, and *T. contortum* (AF013031.1), *T. tenue* (AF013033.1) in clade IV and *T. miru* in clade V (Supplementary Table 4).

We show that *T. nobis* and *T. contortum* are closely related, potentially representing the same non-diazotrophic *Trichodemsium* species. Given that *T. nobis* has not been described morphologically, and the genome of *T. contortum* has not been sequenced, both can be linked through the phylogeny of marker genes. Whereas the previously published genome of *T. nobis* did not contain the *hetR* gene^6^, the clade IV MAG R03 did, and its *hetR* gene sequence was 99.78% similar to that of *T. contortum* (Figure 3; Supplementary Table 4). Complementing our findings, morphological descriptions of *T. contortum* depict the species as single spiral-shaped trichomes, rather than colonies, that are sporadically present in low abundance (< 1% of total *Trichodesmium* biomass) within samples^13,16,53^. MAG R03 lacked the nitrogenase gene cluster, confirming that this clade is non-diazotrophic.

Our results indicate that *hetR* phylotyping can overestimate the diversity within *Trichodesmium* populations, in particular regarding clade II. Intriguingly, both *T. thiebautii* MAGs contained two distinct *hetR* copies (906 bp), which had 28 (MAG R02) and 29 bp (MAG T02) single nucleotide polymorphisms within a sequence. Further inspection of the two copies showed that they were encoded in close vicinity of each other and that the synteny is highly conserved across all the *Trichodesmium* genomes (Figure 4). These genes are likely paralogs of each other^54^ although it is still unclear how *T. thiebautii* benefits from retaining both *hetR* sequences. Each *hetR* sequence was placed into a distinct phylogenetic clade, where one copy grouped with the *hetR* sequence of *T. aureum* (clade II), and the other one was placed within clade I, together with *T. thiebautii*, *T. pelagicum* and *T. hildebrandtii* (Figure 3). Amplicon sequencing of the *hetR* gene showed highly similar abundances of *T. thiebautii* and *T. aureum* in all the samples. Therefore, this is a likely result of *hetR* paralog sequencing within a population of single *T. thiebautii* species (Supplementary Figure 1). Alternatively, *T. thiebautii* and *T. aureum* may represent the morphological diversity within the same genotype or species. We note that the Atlantic *T. thiebautii* contained one *hetR* paralog that clustered with the *hetR* sequence of clade IV (*T. erythraeum*), yet we cannot exclude a binning artifact in this case, as the sequences did not belong to a single scaffold.

**Figure 4.**
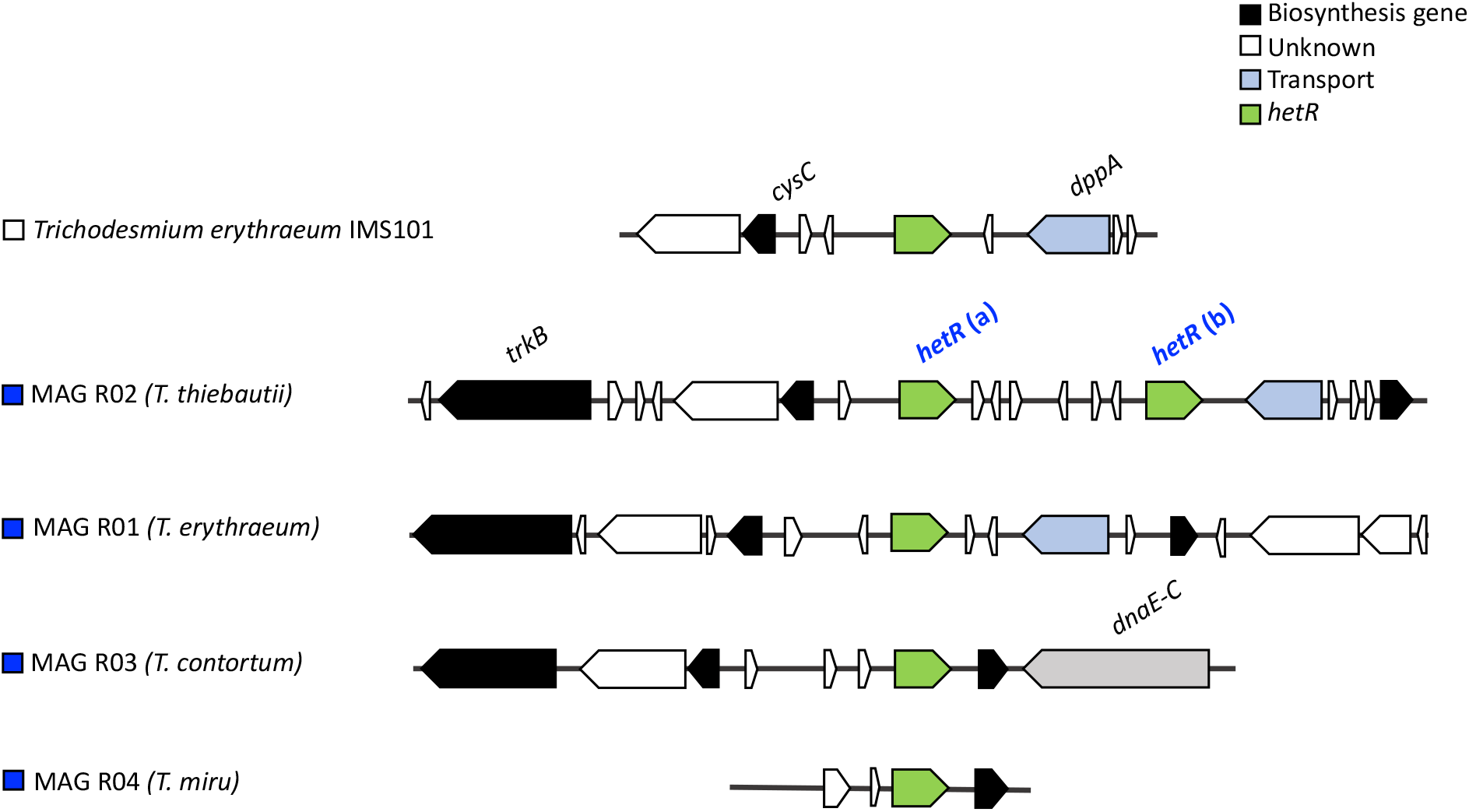
The *hetR* gene clusters in *Trichodesmium* MAGs and *T. erythraeum* IMS101. The synteny of the *hetR* genes (green) and the neighboring genes is conserved. MAG R02 (*T. thiebautii*) contains two *hetR* genes (and are likely paralogs of each other).

### Conclusions and Future Perspectives

Morphotype metagenomics enabled us to better capture the diversity that is present within *Trichodesmium* populations while revisiting past attempts to differentiate between colonies^55,56^. We were able to reconstruct the genomes of four different *Trichodesmium* species isolated from the Red Sea, that were subsequently assigned to four different *Trichodesmium* clades including the more elusive non-diazotrophic clades IV and V. We show that *hetR* gene-based phylotyping could overestimate *Trichodesmium* diversity, highlighting the importance of re-assessing past attempts at phylogeny. Overall, the phylogenetic and functional diversity of *Trichodesmium* in the world’s oceans is still poorly understood to date. Thus, genomics in combination with gene expression and physiology will remain an important avenue for future exploration of these key species.

## Data Availability

The raw sequencing reads from the metagenomic study were deposited at NCBI under the BioProject PRJNA804487. The bioinformatic analysis and scripts were compiled in an R markdown file, together with the genomic FASTA files of the *Trichodesmium* MAGs constructed for this paper, and are publicly available using the following GitHub repository link: https://github.com/cocokoedooder/Trichodesmium_HetR/blob/main/README.md

## Acknowledgments

This work was funded by the Israel Science Foundation (260/21) and the Israel-USA Bi-National Science Foundation (2020041) to YS. This work was also financially supported in part by the Schulich Marine Studies Initiative to IBF. MRB acknowledges the support of the Israel Science Foundation (913/19), the U.S.-Israel Binational Science Foundation (2019055) and the Israel Ministry of Science and Technology (1126). FZ thanks the PBC Fellowship Program for Outstanding Chinese and Indian Post-Doctoral Fellows. We would like to thank Murielle Dray for technical support.

## Competing Interests

The authors of this paper declare no conflict of interest.

## Author Contributions

CK, YS, IB-F and MR-B conceived the study. YS and IB-F acquired financial support. CK, FZ, SB collected *Trichodesmium* colonies and extracted DNA for metagenomics. CK analyzed the metagenomic data under the guidance of MRB. EL performed the seasonal *hetR* amplicon sequencing and data analysis under the guidance of IB-F. CK, EL and MR-B wrote the manuscript with contributions from other co-authors. All authors discussed the results and revised the final version of the manuscript.

## Supplementary Information

**Supplementary Figure 1.**
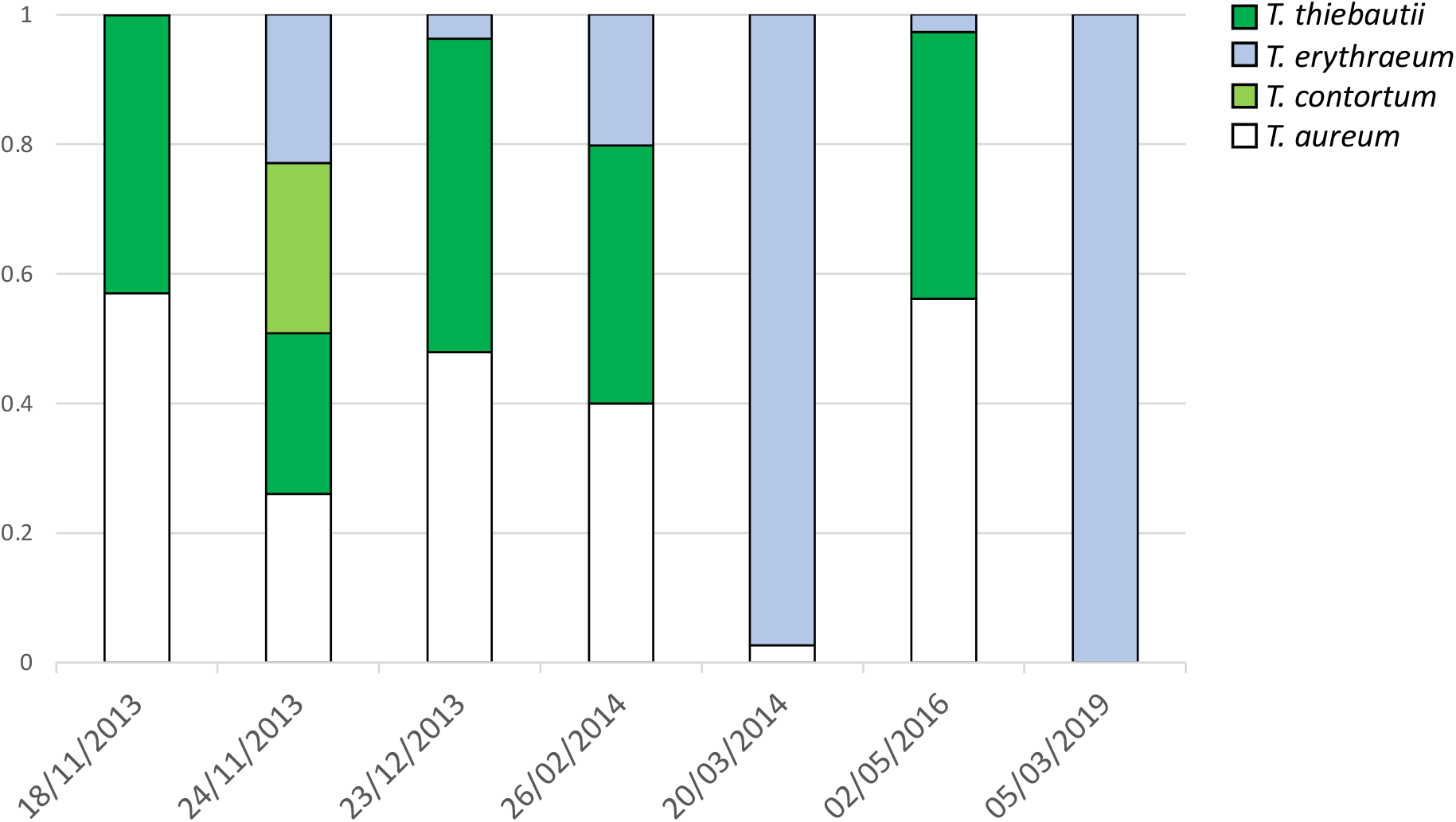
Displaying the relative abundance of *hetR* sequences from (puff and tuft) *Trichodesmium* colonies isolated from the Red Sea. Note that *T. thiebautii* and *T. aureum* are in similar proportions to each other. The presence of *T. aureum* is likely an artifact and instead represents a paralog *hetR* sequence present in *T. thiebautii*, thereby effectively over-estimating *Trichodesmium* diversity within the sample.

**Supplementary Table 1.** General information about the *Trichodesmium* MAGs derived from this study.

| MAG ID | Accession Number | HetR phylotype | Size (Mbp) | Completeness | Redundancy | A. Dense-puff (%) | B. Long-puff (%) | C. Tuft (%) |
| --- | --- | --- | --- | --- | --- | --- | --- | --- |
| MAG R01 | SAMN25809967 | *T. erythraeum* | 6.641 | 95.8 | 0.3 | 2.8 | 4.4 | 83 |
| MAG R02 | SAMN25809968 | *T. thiebautii*/ *T. aureum* | 6.769 | 94.8 | 0.4 | 77.1 | 78.1 | 4.3 |
| MAG R03 | SAMN25809969 | *T. nobis* | 5.841 | 94.0 | 2.5 | 5 | 8 | 3.1 |
| MAG R04 | SAMN25809970 | *T. contortum* | 4.176 | 83.0 | 3.1 | 0.6 | 1.3 | 0.6 |

## Notes

https://github.com/cocokoedooder/Trichodesmium_HetR

### Competing Interest Statement

The authors have declared no competing interest.

